# Subendothelial Calcification Increases Surface Roughness and Promotes Lipid Deposition into the Arterial Wall

**DOI:** 10.1101/2024.09.30.615966

**Authors:** Nina D. Kosciuszek, David Petrosian, Navya Voleti, Param Dave, Ian Kelly, Daniel Moussouros, Josef Davidov, Anton Mararenko, Kelly A. Borges, Saud A. Nasruddin, Mugdha V. Padalkar, Maria M. Plummer, Jose Luis Millan, Dorinamaria Carka, Brian L. Beatty, Olga V. Savinova

**Author notes:** Correspondence: Brian L. Beatty, Department of Anatomy, New York Institute of Technology College of Osteopathic Medicine, PO box 8000, Northern Blvd, Old Westbury, NY 11568,; Olga V. Savinova, Department of Biomedical Sciences, New York Institute of Technology College of Osteopathic Medicine, PO box 8000, Northern Blvd, Old Westbury, NY 11568.

## Abstract

**Objective:** We have previously demonstrated that subendothelial calcification accelerates atherosclerosis in mice. This study addresses a mechanism by which subendothelial calcifications can increase low-density lipoprotein (LDL) uptake into the arterial wall.

**Methods:** Mice overexpressing tissue-nonspecific alkaline phosphatase (TNAP) in endothelial cells (eTNAP mice) were used as a model of calcification. Calcification and atherosclerosis were detected by micro-computed tomography (micro-CT) and histology. The endothelial roughness was characterized by surface metrology. A fluid-structure interaction model was used to calculate wall shear stress (WSS). The uptake of fluorescent LDL was traced *in vitro* and *in vivo*. Human arteries were assessed for the prevalence of internal elastic lamina (IEL) calcification.

**Results:** eTNAP mice developed more severe aortic atherosclerosis than controls on the LDL receptor mutant background (p<0.01). Subendothelial calcifications in eTNAP mice were confirmed by micro-CT. An increase in aortic surface area roughness, including the height, volume, and steepness parameters, was observed in eTNAP mice compared to controls (p<0.01). Calcifications affected near-wall hemodynamics, creating pockets of reduced WSS. Endothelial cells cultured on rough surfaces showed increased LDL uptake compared to cells cultured on smooth collagen (p<0.0001). Fluorescent LDLs were traced to subendothelial calcifications in eTNAP mice but not in controls. In humans, IEL calcification was prevalent in older adults and inversely correlated with arterial diameter (p<0.05).

**Conclusion:** Subendothelial calcification is sufficient to perturb near-wall hemodynamics, creating localized areas of reduced WSS, consistent with increased LDL uptake near calcified lesions. Subendothelial calcification may represent an alternative or concurrent mechanism for the initiation of atherosclerosis.

**Research perspective:** *What is new?:*

- We tested a novel hypothesis that subendothelial microcalcification can initiate atherosclerosis.
- The study demonstrated that micron-sized subendothelial calcifications, induced by the overexpression of tissue-nonspecific alkaline phosphatase in the endothelium, are sufficient to perturb local hemodynamics, creating pockets of low wall shear stress, consistent with an increase in low-density lipoprotein uptake and deposition into the arterial wall in juxtaposition to calcified lesions.

*What are the clinical implications?:*

- We suggest that calcification of internal elastic lamina of medium-sized arteries may represent early lesions initiating atherosclerosis; however, the epidemiologic evidence for this theory is currently unavailable.

## Introduction

Atherosclerotic cardiovascular disease (ASCVD) is the leading cause of death worldwide.^1^ Many risk factors contribute to the development and progression of ASCVD, including elevated plasma cholesterol, hyperglycemia, and hypertension. These modifiable risk factors are targeted therapeutically to improve ASCVD outcomes.^2^ Vascular calcification is another risk factor associated with ASCVD.^3^ Coronary artery calcium (CAC) score independently predicts major adverse cardiovascular events (MACE).^4–8^ Yet, it is unknown whether vascular calcification directly contributes to the pathogenesis of ASCVD. A basic understanding of the interaction between calcification and atherosclerosis could position vascular calcification as a modifiable risk factor that can be targeted therapeutically to mitigate ASCVD.

Vascular calcification is a regulated process, the mechanism of which partly resembles that of skeletal biomineralization.^9^ Tissue-nonspecific alkaline phosphatase (TNAP) is an enzyme essential for skeletal biomineralization.^10^ TNAP creates mineralization-permissive conditions by hydrolyzing inorganic pyrophosphate (PPi), a potent endogenous inhibitor of biomineralization, and establishing the appropriate PPi to inorganic phosphate (Pi) ratio.^11^ Increased TNAP activity is detected in calcified arterial lesions in experimental animals and humans, underscoring its role in the pathologic biomineralization of arterial tissues.^12–15^

Arterial calcification can present in many forms and affect different arterial layers. It has been found that microcalcification in atherosclerotic plaques can cause a lesion to become unstable, increasing the risk of plaque rupture.^16, 17^ In addition to spotty microcalcifications, calcified nodules and superficial sheets can render atherosclerotic plaques susceptible to erosion and subsequent thrombosis and a vessel occlusion.^18, 19^ Calcifications formed in the subendothelial space, such as those nucleated in the internal elastic lamina (IEL), can encroach on the adjacent intima, altering the topology of endothelial surface.^20^ Endothelial cells respond to blood flow, and these reactions can be either atheroprotective or promote atherosclerosis.^21^ The fluid drag exerted by blood flow on the endothelial surface is characterized by the magnitude and direction of wall shear stress (WSS). Low and oscillatory WSS can lead to atherosclerosis.^21^ Limited studies have evaluated the effect of minor irregularities of luminal surface on WSS^22, 23^ We hypothesized that TNAP-induced subendothelial calcification (1) is associated with increased surface roughness, (2) contributes to pathologic redistribution of WSS, and (3) increases low-density lipoprotein (LDL) uptake. This pathologic sequence could explain our observation of increased atherosclerosis burden in mice with TNAP-induced subendothelial calcification.

## Methods

### Animal studies

Animal studies were approved by the Institutional Animal Care and Use Committees (IACUC) of Sanford Research (Sioux Falls, SD) and the New York Institute of Technology College of Osteopathic Medicine (NYITCOM, Old Westbury, NY) and complied with the National Institutes of Health guidelines for humane treatment of laboratory animals. Mice were euthanized by exsanguination under 5% isoflurane in oxygen or by carbon dioxide inhalation.

Two independent cohorts of mice with a point mutation in the LDL receptor gene (*ldlr^whc/whc^*, abbreviated WHC for wicked high cholesterol^24^) were produced on a mixed (129:B6) genetic background in two animal facilities, the Sanford Research between 2014-2015 (cohort 1, Sioufx Falls, SD) and NYITCOM in 2016-2017 (cohort 2, Old Westbury, NY). The assessment of the aortic root and coronary atherosclerosis in these mice was previously reported.^25^ Whole-body formalin-perfused specimens were stored in formalin until further experiments in 2016 (cohort 1, 13 mice) and 2018 (cohort 2, 11 mice). Each cohort consisted of a control group (*ldlr^whc/whc^*, WHC only) and an endothelial TNAP overexpressor group (WHC-eTNAP). Both cohorts were fed an atherogenic diet (1.25% cholesterol, Harlan/Envigo TD.02028) for eight weeks. All mice were males because of a limitation imposed by their X chromosome-linked TNAP transgene^26^ that affected cardiovascular fitness, limiting breeding performance. eTNAP mice without the *Ldlr* mutation were backcrossed to a C57BL/6 genetic background for eight generations at NYITCOM. These mice and their control littermates were used for micro-computed tomography (micro-CT) and *in vivo* fluorescent LDL uptake experiments. Data were collected by researchers who were blinded to the animal genotypes.

### Assessment of calcification and atherosclerosis in animal models

Calcification and atherosclerosis were assessed by calcium-specific alizarin red and oil red O staining, specific for lipids, of flat-mounted preparations of the thoracic aorta and quantified by measuring the affected areas using the ImageJ software. In addition, calcification and lipid deposition were qualitatively assessed in cross-sections of the thoracic aortas. These sections were counterstained with hematoxylin and examined with light microscopy.

### Blood chemistry

Lithium heparin plasma was isolated from mice in terminal experiments after a five-hour fast. Plasma total cholesterol and triglycerides were measured using clinical chemistry reagents (Pointe Scientific, Canton, MI).

### Micro-CT

Immediately after sacrifice, a subset of eTNAP mice on the C57BL/6 genetic background was perfused with 500 IU of heparin in saline, followed by MICROFIL MV-122 contrast polymer injection (Microfil, Boulder, CO) *via* the left ventricle (n = 3/group). Tissues were fixed in formalin, and the abdominal aortas were dissected with the spine. The specimens were scanned using a SkyScan 1173 micro-CT scanner (Bruker, Billerica, MA) at 5.7-micron resolution. The resulting images were reconstructed in NRecon (Bruker) and segmented in Dragonfly ORS (Object Research Systems, Montreal, Canada). Five percent of data were used to train a UNet5 Deep Learning model (in Dragonfly) that segmented the lumen, wall, and calcifications within the aorta and branching vessels.

### Surface metrology

Flat-mounted preparations of suprarenal abdominal aortas were used for surface metrology. These aorta segments were dissected from formalin-fixed whole-body preparations, opened longitudinally, mounted on slides using a permanent mounting medium (Clearium, Electron Microscopy Sciences), and air-dried. The luminal surfaces were scanned using a Sensofar S Neox optical white light reflectance confocal microscope (Terrassa, Spain) at 1-micron vertical resolution. Each sample, measuring 0.5 x 0.7 mm, was analyzed using the SensoMAP software (Terrassa, Spain) to determine their areal roughness parameters (ISO25178-2) as described previously.^27^ Files were subjected to the form removal SensoVIEW’s operator to remove the tilt and unintentional effects of warped surfaces (polynomial-3 function). Other surface artifacts were removed with the retouching function. Samples that contained areal data gaps due to the excessive steepness or retouching were subjected to the filling of non-measured points, which replaced data gaps with a smooth surface formed from the surrounding points. A higher vertical resolution scan was obtained with a 50X objective at 0.2 microns vertical resolution to reconstruct surface geometry for computational modeling.

### Fluid-structure interaction modeling

We previously reported the details of our Fluid-Structure Interaction (FSI) modeling at the CMBBE (Computer Methods in Biomechanics and Biomedical Engineering) Symposium.^28^ Briefly, to construct the realistic geometry of an eTNAP mouse aorta affected by subendothelial calcification, we used the high-resolution optical scan of the thoracic aorta. The computational mesh points of the luminal surface were mapped onto a cylindrical geometry. This geometry was imported in MatLab and used for 3D modeling by the Non-Uniform Rational B-Splines (NURBS) method.^29^ The resulting geometry was then imported in COMSOL Multiphysics 5.4, and a fully coupled FSI model was constructed as a periodic 55-degree slice of a 2 mm segment of the aorta with a radius of 0.6 mm. The endothelium was considered 3.8 microns thick and rigidly attached to the arterial wall. The endothelial layer was modeled as hyperelastic (Neo-Hookean) material with Young’s modulus of 2 kPa.^30^ Calcified lesions were modeled as linear elastic material with five times the Young’s modulus of the endothelial layer. A fully formed parabolic laminar flow with a mean velocity of 17 cm/s was applied as an inlet boundary condition, and a zero-pressure boundary was imposed on the outlet. Blood was modeled as a Newtonian fluid with a constant viscosity of 0.00282 Pa·s. The Navier-Stokes equations were solved in the fluid domain, and Newton’s equation on the calcified lesions and endothelium domains.

### *In vitro* LDL uptake

Human umbilical cord endothelial cells (HUVEC) were obtained from the American Type Culture Collection (ATCC, Manassas, WA) and cultured in F-12K Medium supplemented with 10% Fetal Bovine Serum (FBS), 0.1 mg/mL heparin and 30 µg/mL Endothelial Cell Growth Supplement (ECGS, Cell Applications, San Diego, CA). The experiments were conducted in 24-well plates coated with 2% gelatin (10 µl/cm^2^) with or without hydroxyapatite beads (3 mg/ml, ∼2000 particles/cm^2^, average diameter 10 microns, cat # 900203, MilliporeSigma, St. Louse, MO).

Hydroxyapatite was used to model subendothelial calcifications. We used an orbital shaker operated at 75 rpm inside an incubator to model constant fluid flow over the surface. Endothelial cells were plated at 2 x 10^5^ cells per well and allowed to adhere overnight under static conditions. Human acetylated (Ac), oxidized (Ox), and native LDL labeled with DiI fluorescent dye (1,1′-Dioctadecyl-3,3,3′,3′-tetramethylindocarbocyanine perchlorate) were obtained from Kalen (Balen Biochemical, Montgomery Village, MD) and used in all experiments at 30 µg/ml concentration. Each condition and time-point were run in duplicate, and LDL uptake experiments were repeated two times. The amounts of fluorescent LDL uptake were measured by flow cytometry after labeling of cells with Invitrogen LIVE/DEAD® Fixable Far-Red stain (ThermoFisher Scientific, Waltham, MA) and formalin fixation. ACCURI C6 fluorescent flow analyzer (BD Biosciences, San Jose, CA) was used for these studies. Up to 10,000 cell scatter-gated events were collected from each well and further gated to exclude dead cells. Fluorescent intensity was plotted against time, and one-phase association curves were fitted using GraphPad Prism 9 software (GraphPad, La Jolla, California) to determine the maximum LDL uptake.

### *In vivo* LDL uptake

16-week-old eTNAP and control mice on the C57BL/6 genetic background (n = 3 per group) were injected with 75 µg (150 µl) of DiI-labeled native LDL (Kalen) via the retroorbital sinus. Mice were sacrificed 6 hours after the LDL injection and perfused with formalin. Aortic arches were dissected, embedded in OCT, and sectioned longitudinally. Mesenteries were dissected, and cryo-sections were prepared to visualize cross-sections of mesenteric arteries. The sections were cover-slipped in Fluoromount-G™ Slide Mounting Medium with DAPI (4′,6- diamidino-2-phenylindole; Electron Microscopy Sciences, Hatfield, PA) and examined under a brightfield microscope to identify irregularities in the luminal surface in eTNAP mice. 5 to 6 lesions per sample were identified and imaged under a fluorescent microscope with red (DiI) and blue (DAPI) emission filters. The control samples were similarly imaged at the anatomically equivalent positions along the aortic arch and in the equally sized mesenteric arteries.

### Human tissues

Using cadaveric tissues, we investigated the prevalence of calcification in the internal elastic lamina (IEL), tunica media, and intima in human arteries. Individuals who have chosen to donate their bodies after death consented to allow their tissues to be used for medical research. The tissues were de-identified and did not require an IRB approval. The demograthic data available in the donors included sex, age, and the cause of death, as stated on the death certificates. Non-decalcified tissues were cryo-sectioned and analyzed using alizarin red staining for calcium. We recorded the prevalence of any calcification in the intimal, IEL, and media layers of the common carotid arterys (CCA, 2 cm proximal of the carotid bulb), internal and external carotid arteries (ICA-C1 segment and ECA, 1 cm distal from the carotid bulb); intracranial segment of ICA (ICA-C7); middle cerebral arteries M1 segment (MCA, 0.5 cm proximal and distal of the circle of Willis, respectively); and the marginal artery of the colon (Drummond marginal artery, DMA) sampled at splenic flexure.

### Statistical Analysis

Statistical analyses were performed with GraphPad Prism 9 (GraphPad, La Jolla, California). Data distribution was tested using the D’Agostino-Pearson normality test. Non-normally distributed data were log-transformed before the analysis. If log-transformed data did not meet the normality assumptions, non-parametric tests were used to compare variables. Two groups were compared using an unpaired Student’s t-test or a Mann-Whitney test. The curve-fitting of a one-phase association equation (GraphPad Prism 9) was performed to quantify the plateau phases of *in vitro* LDL uptake, which were pair-wise compared (rough *vs.* smooth surfaces) by an extra sum-of-squares F test. In two independent trials, the reproducibility of LDL uptake was assessed by a two-way ANOVA to identify the effects of surface roughness on LDL uptake plateaus. Linear regression analysis was used to test the association between vessel diameter and the prevalence of calcification in different vascular layers (intima, media, and IEL) in human tissues. Normally distributed variables are presented as mean ± standard deviation, and non-normally distributed variables as median and 95% confidence intervals. A value of p < 0.05 was accepted as statistically significant.

## Results

### TNAP-induced subendothelial calcification accelerates aortic atherosclerosis

We hypothesized that small, calcified subendothelial lesions would be sufficient to increase LDL deposition into the arterial wall at the sites of calcification due to their effects on the near-wall arterial hemodynamics. We first confirmed that eTNAP mice on the *ldlr* mutant (129:B6 mixed) genetic background develop aortic calcifications and accelerated atherosclerosis consistent with our previous findings.^25^ *En-face* preparations of the thoracic aortas from the first cohort of 16- week-old eTNAP and control mice were divided between alizarin red and oil red O staining to quantify aortic calcium and lipid deposition. Compared to controls, eTNAP mice showed a larger area affected by calcifications (4.0 ± 2.5% *vs.* 0.7 ± 1.5%, p < 0.05, Figure 1A) and more extensive aortic atherosclerosis (28.1 ± 8.3% *vs.* 8.6 ± 4.3, p < 0.01, Figure 1B). The cross-sections of thoracic aortas from the second cohort of mice were assessed by alizarin red and oil red O with hematoxylin staining to detect calcification and lipid deposition (Figure 1C, D). We noted a co-localization of lipids (oil red O-positive) with calcifications (stained deep blue by hematoxylin) in the aortic wall of eTNAP mice but not in control mice (Figure 1D). At the same time, the plasma lipids (total cholesterol and triglycerides) were not different between eTNAP and control mice (Figure 1E, F). A high-resolution (5.7 microns) contrast-enhanced micro-CT of the abdominal aortas confirmed the presence of calcified lesions in the eTNAP mice on a regular C57BL/6 genetic background but not in control mice (Figure 1G, H), as was it was previously demonstrated on the 129:B6.^31^

**Figure 1.**
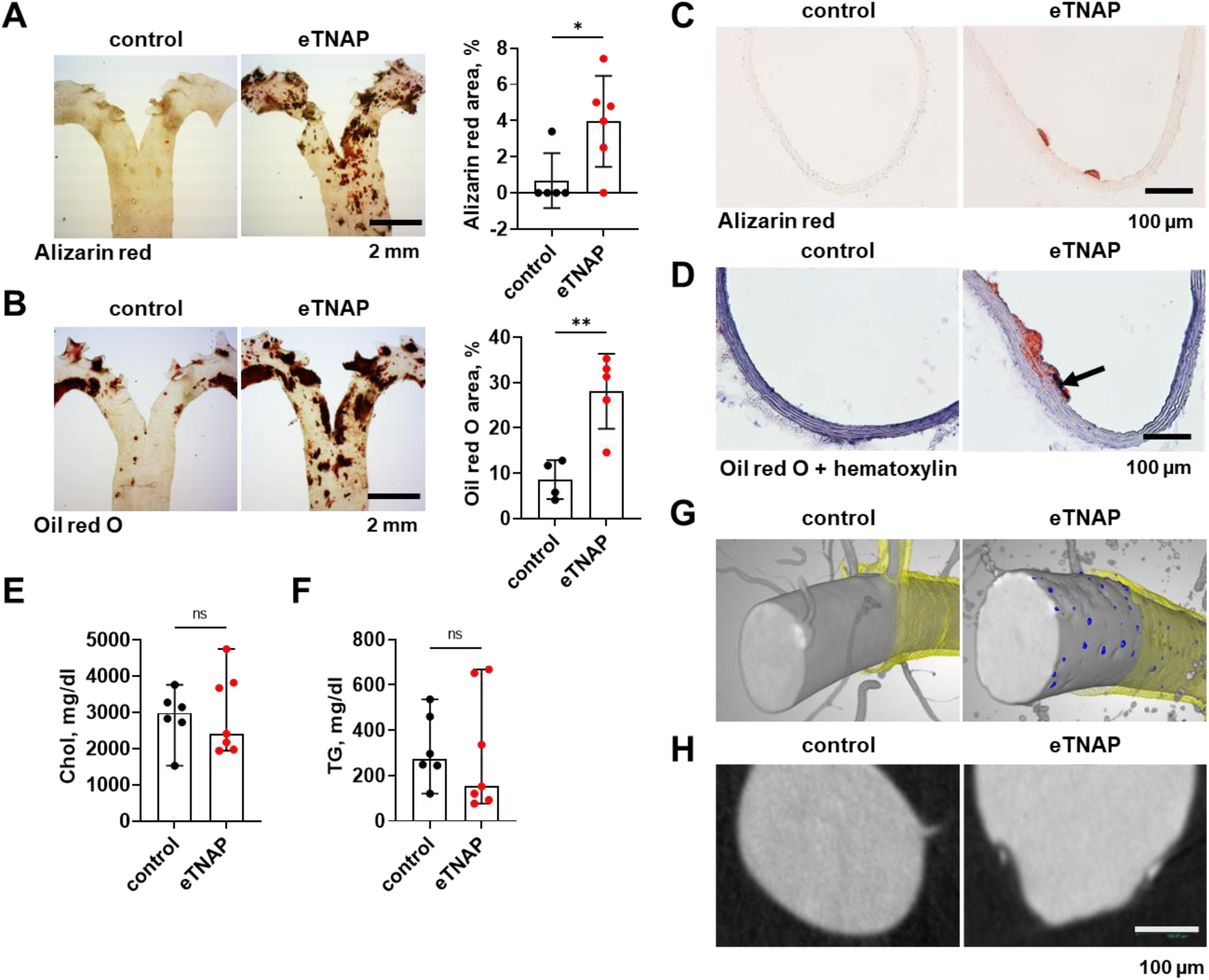
Subendothelial calcification and atherosclerosis in the aorta of eTNAP and control mice. A. Representative images and quantification of alizarin red-positive staining in the thoracic aortas, n = 5-6/group; B. Representative images and quantification of oil red O, n = 4-5/group; C. Detection of subendothelial calcification by alizarin red staining; D. Representative images of oil red O staining with hematoxylin; arrow points to a site of calcification stained deep blue; E. Plasma total cholesterol, n = 6-7/group; E. Plasma triglycerides, n = 6-7/group; (A-E) mice in the *ldlr* mutant background G. 3D reconstruction of the abdominal aorta from eTNAP and control by micro-CT; segmentation of the arterial wall (yellow) and calcified nodules (blue); H. Representative micro-CT slices of eTNAP and control aortas. (G,H) mice of the C57BL/6 genetic background; *, p < 0.05; **, p < 0.01; ns, not significant.

### Subendothelial calcification increases endothelial surface roughness

We used surface metrology, a precise method, to quantitatively describe and compare the luminal surfaces of flat-mounted preparation of abdominal aortas from the first cohort of eTNAP and control mice (Figure 2A). We found that, compared to control aortas, the surface of the aorta in eTNAP mice was characterized by increased maximum peak height (Sp, p < 0.01, Figure 2B), peak material volume (Vmp, p < 0.01, Figure 2C), steepness/roundness of surface features (root-mean-square gradient, Sdq, p < 0.01), and surface area contributed by texture (developed interfacial areal ratio, Sdr, p < 0.05; Figure 2E). The experiment was repeated on the second cohort of mice, and the data from both experiments were analyzed by a two-way ANOVA. After adjusting for the trial number (a source of technical and biologic variability), the two-way ANOVA showed significant effects of endothelial TNAP overexpression on multiple areal roughness parameters, demonstrating an increase in the aortic luminal surface roughness in eTNAP mice compared to controls (Table 1).

**Figure 2.**
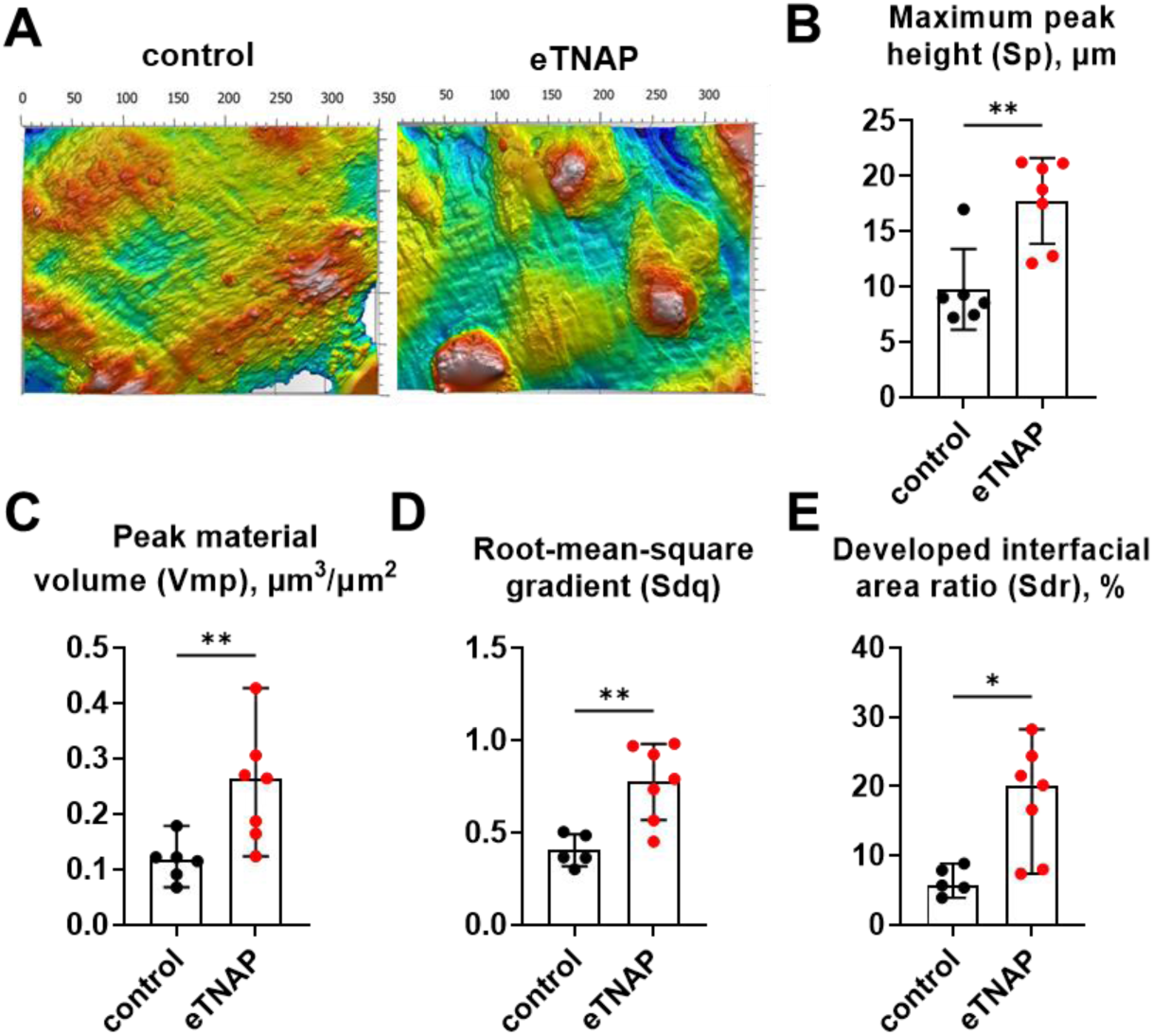
Effects of subendothelial calcification on areal surface parameters (ISO 25178-2) in eTNAP and control aorta. A. Representative plots of abdominal aortas obtained by surface metrology; B. Maximum peak height (Sp); C. Peak material volume (Vmp); D. Root-mean- square gradient (Sdq); E. Developed interfacial area ratio (Sdr). All mice were on the *ldlr* mutant background *, p < 0.05; **, p < 0.01; n = 5-7/group.

**Table 1.**
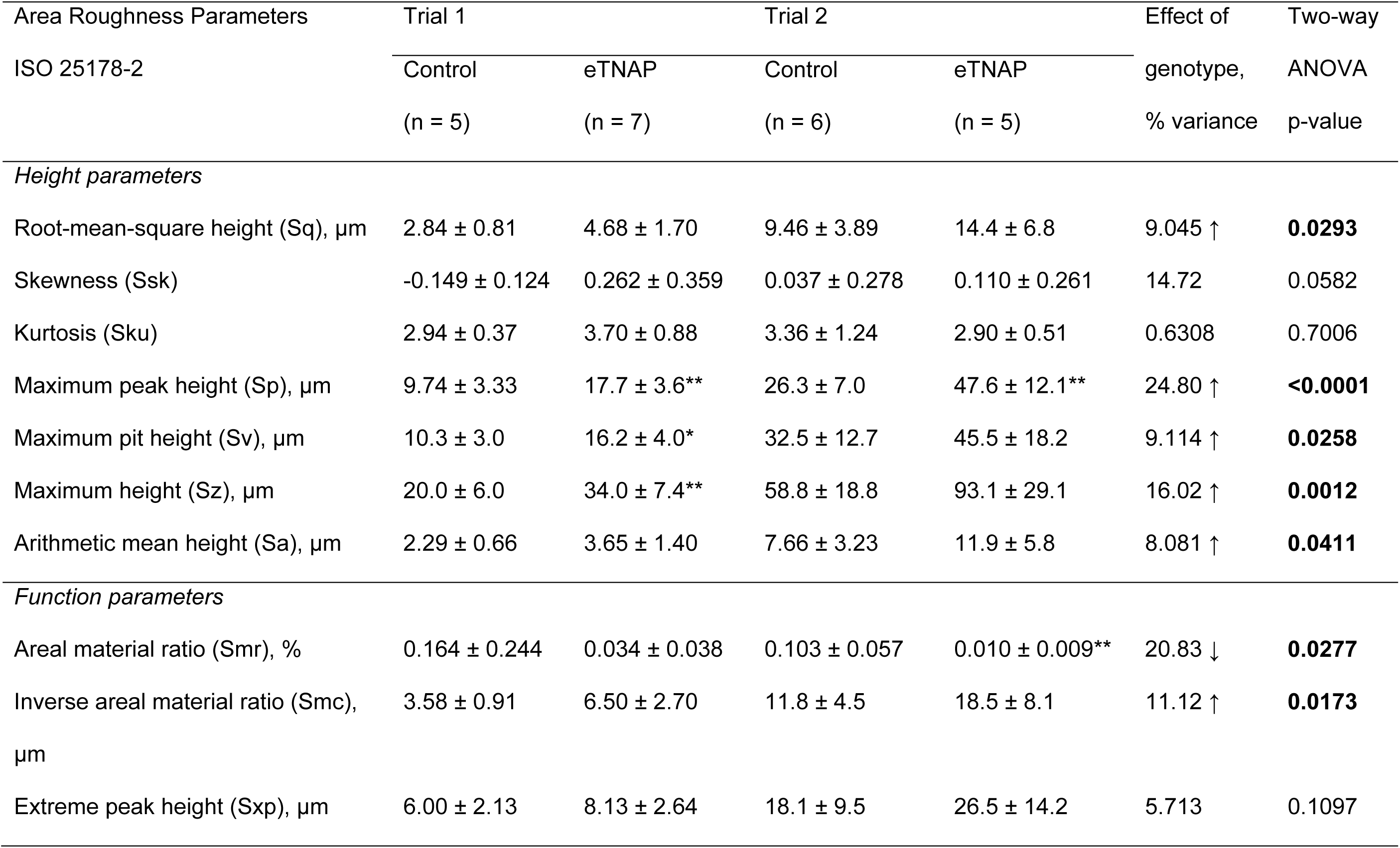
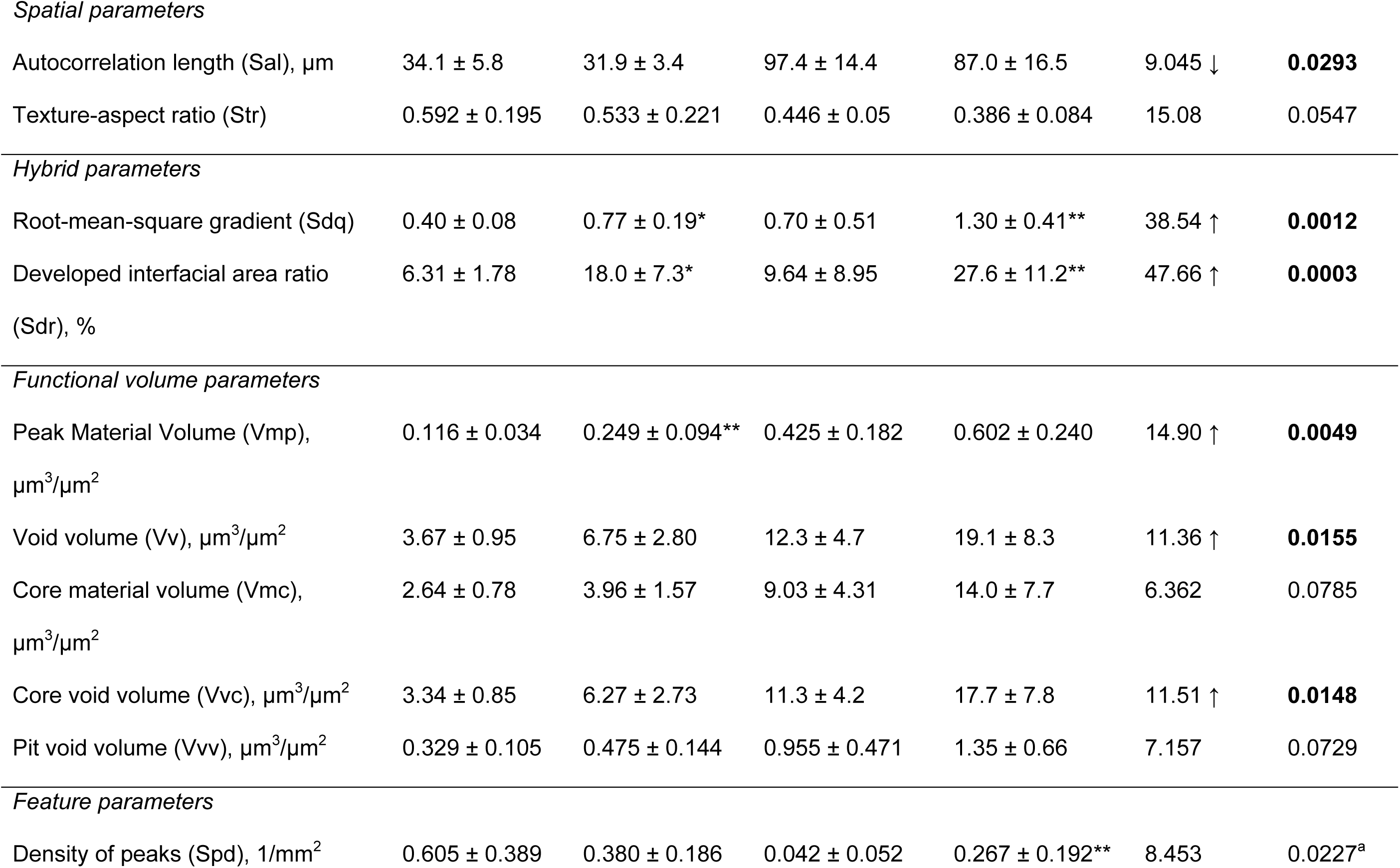
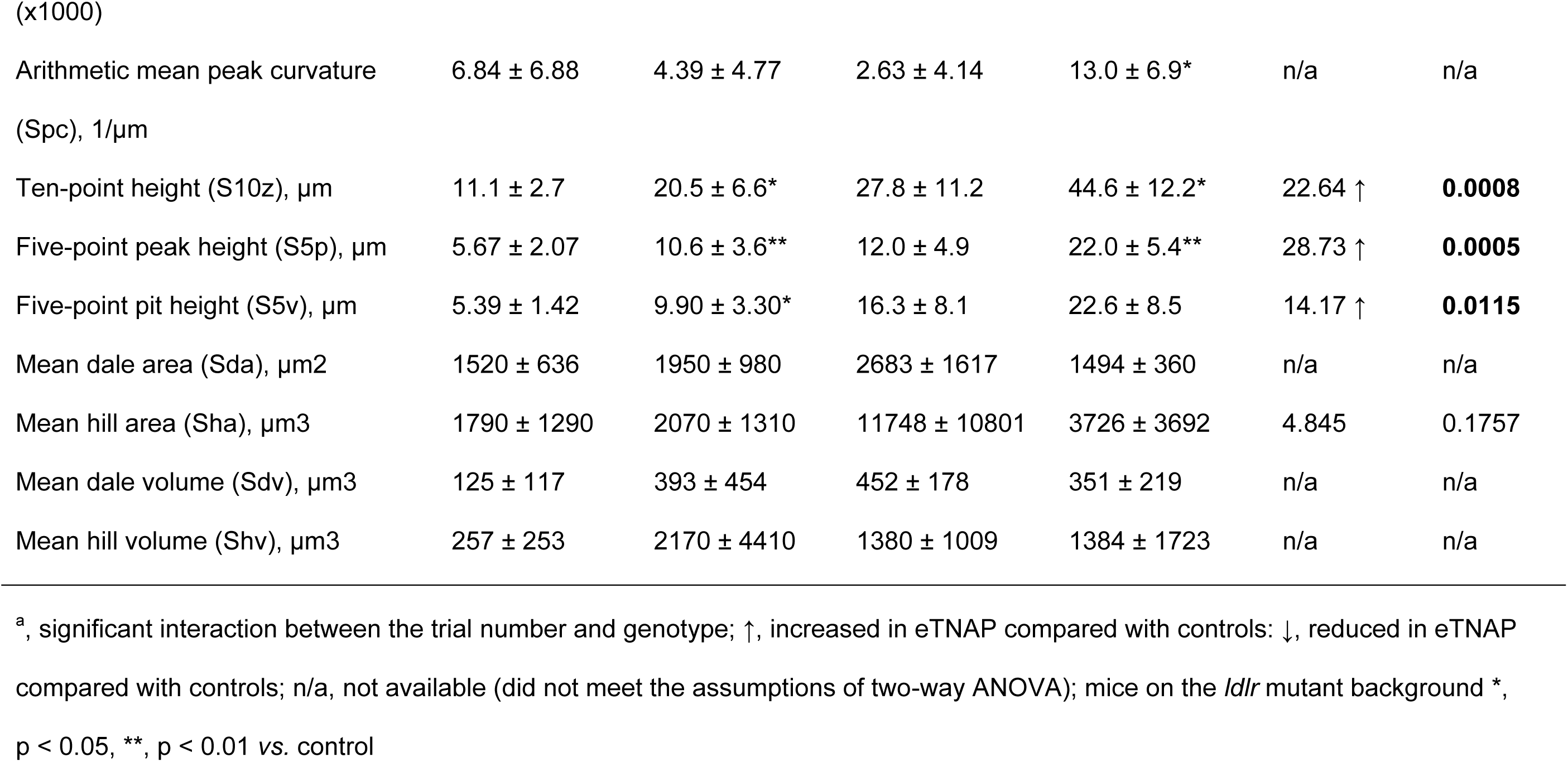
Area roughness parameters of abdominal aortas from two cohorts of eTANP and control mice, assessed by surface metrology and analyzed by two-way ANOVA accounting for the effects of trial number and eTNAP genotype.

### Aberrant WSS distribution in the aorta of eTNAP mice

We hypothesized that the relationship between increased endothelial surface roughness and accelerated atherosclerosis in eTNAP mice could be explained by the effect of surface roughness on wall shear stress (WSS). To test this hypothesis, we constructed an FSI model. Stitched optical confocal scans of a representative eTNAP mouse aorta collected at 0.2-micron resolution were used to reconstruct the geometry of the luminal surface of the thoracic aorta (Figure 3A). The surface was optimized for geometric import and computation. The details of computational procedures were previously reported at the Computer Methods in Biomechanics and Biomedical Engineering Symposium.^28^ The distribution of the WSS magnitude demonstrated a 30-50% reduction of WSS near standalone small lesions compared to smooth regions of the vessel, with the largest reduction observed behind the lesions relative to the blood flow direction. The regions with congregated lesions with irregular boundaries showed a more complex WSS distribution, with the magnitude of WSS reduced to zero at some spots (Figure 3B).

**Figure 3.**
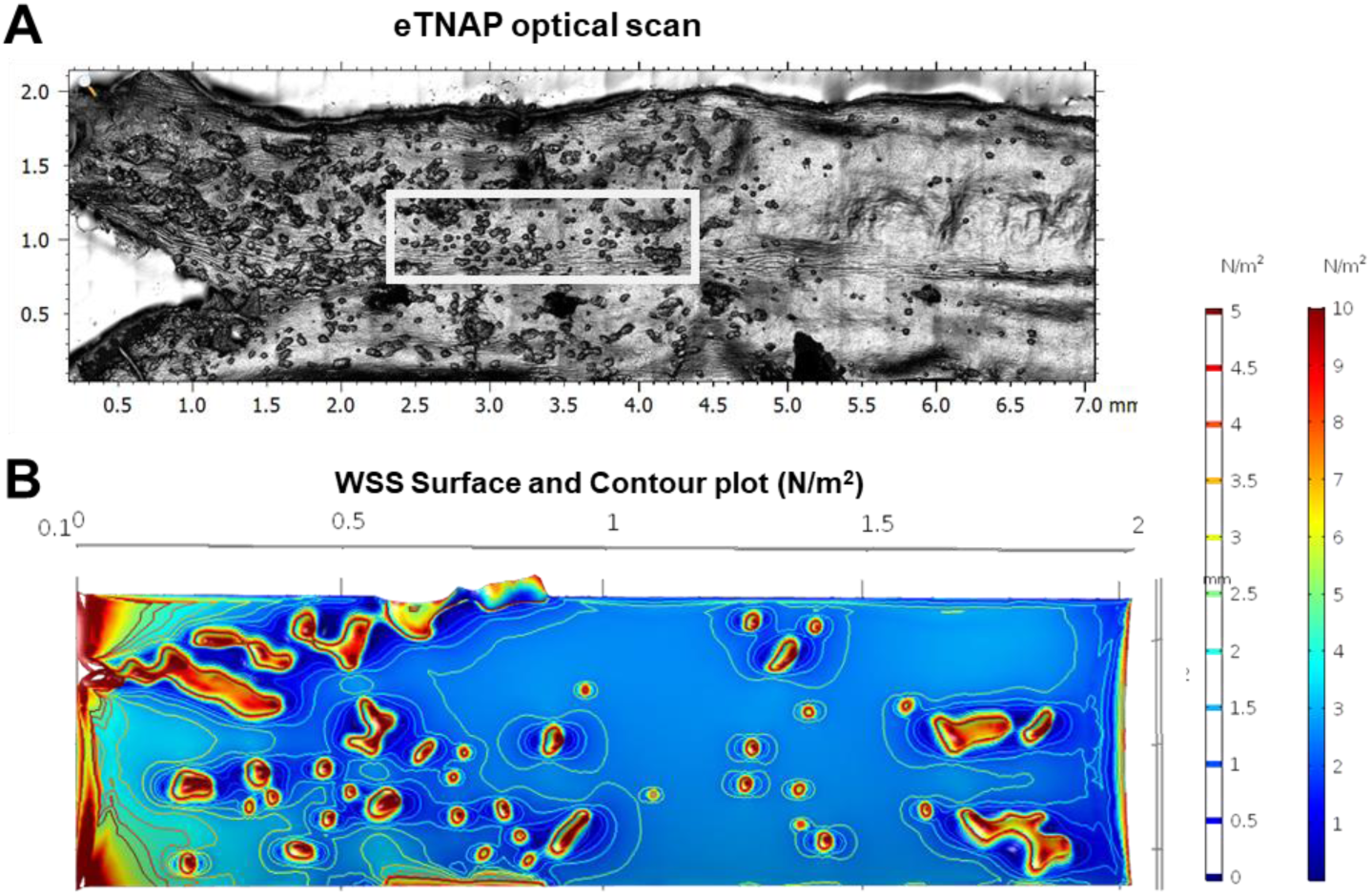
Effect of increased surface roughness on the WSS distribution in the aorta of eTNAP mice on the C57BL/6 genetic background: FSI modeling. A. Surface plot of the thoracic aorta from eTNAP mouse scanned at 0.2-micron resolution; the boxed area represents the surface used to construct realistic geometry for the FSI model; B. WSS magnitude contour and surface plot on the fluid-structure interface.

### Comparison of LDL uptake by endothelial cells cultured on smooth and rough surfaces

Next, we tested our central hypothesis that increased surface roughness promotes LDL uptake. We first conducted *in vitro* experiments with human endothelial cells (HUVEC). Cell culture plates were coated with gelatin (G) or gelatin mixed with hydroxyapatite particles of an average diameter of 10 microns (G+HA) to model smooth and rough surfaces. We cultured HUVEC on a rotating platform inside an incubator at 75 pm to simulate blood flow. In two independent trials, cells were treated with DiI-labeled acetylated (Ac-), oxidized LDL (Ox-), or native LDL at a concentration of 30 µg/ml (Figure 4A). LDL uptake after 0, 2, 4, 8-10, and 24-26 hours was measured by flow cytometry. The average DiI fluorescent intensity of live cells (70-95%) was plotted against time. The plateaus of the one-phase association curves, fitted to the DiI intensity, were taken as maximum LDL uptake under each condition. No differences were found in the Ac- LDL or Ox-LDL uptake between cells cultured on smooth or rough surfaces (Figure 4B, C). The native LDL uptake was increased by 38-52% in cells cultured on rough surfaces compared with cells on smooth surfaces (p < 0.0001, Figure 4D). The LDL uptake data from two trials were then analyzed by a two-way ANOVA, accounting for the trial number (technical/biologic variability) and surface roughness. We found no overall effect of surface roughness on the Ac- LDL or Ox-LDL uptake (Figure 4E, F) but a significant effect on the native LDL uptake (p = 0.0028, Figure 4G).

**Figure 4.**
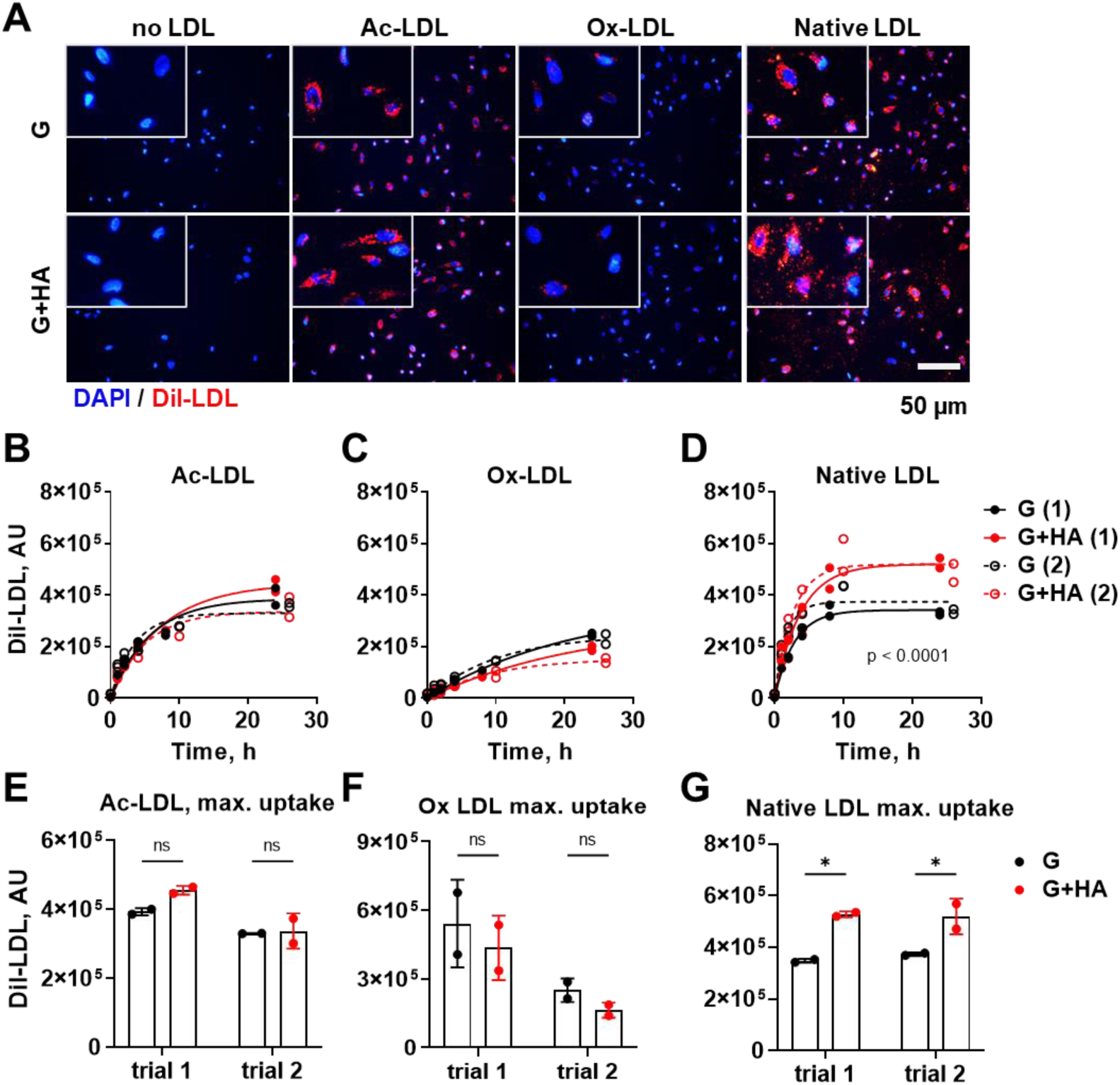
Increased surface roughness promotes native LDL uptake by endothelial cells *in vitro*. A. Representative images (with enlarged inserts, 4x) of human endothelial (HUVEC) cells cultured on smooth (gelatin-coated, G) or rough (gelatin with hydroxyapatite beads, G+HA) surfaces and treated with fluorescently (DiI)-labeled acetylated (Ac-), oxidized (Ox-), or native LDL (red) for four hours, counterstained with DAPI (blue); B-D. One-phase association curves of (DiI)-labeled acetylated (Ac-), oxidized (Ox-), and native LDL with endothelial cells determined by flow cytometry at multiple time points; E-G. Plateaus of the one-phase association LDL uptake curves representing maximum LDL uptake in two trials. *, p < 0.05; ns, not significant.

### The effect of subendothelial calcification on LDL uptake in vivo

Our previous studies have shown that eTNAP mice on the *ldlr* mutant background develop accelerated atherosclerosis.^25^ In this experiment, we used eTNAP mice on a standard C57BL/6 genetic background unaffected by *ldlr* mutations (Figure 1G; micro-CT of subendothelial calcification). Using these eTNAP mice and their control littermates, we wanted to test whether pre-existing subendothelial calcification can trigger LDL uptake at the sites of calcification. 16- week-old eTNAP and control mice (n = 3 per group), on a regular diet, were injected with 75 µg of DiI-labeled native LDL (i.v; 150 μl per mouse). Mice were sacrificed 6 hours after LDL injection and perfused with formalin. The uptake of native DiI-LDL was traced by histology with fluorescent detection of DiI-labeled LDL. The aortic arches were sectioned longitudinally (Figure 5A). The luminal surface was examined for irregularities representing subendothelial calcification. We observed minimal incorporation of DiI-LDL in the intima of the aortic arch of control mice (Figure 5A-i) but an appreciable uptake of DiI-LDL in the aortic arch of eTNAP mice primarily at the sites affected by calcification (Figure 5A-ii). Moreover, we found no incorporation of DiI-LDL in the mesenteric arteries of control mice (Figure 5B-i) but extensive incorporation of DiI-LDL in the intima of calcified mesenteric arteries of eTNAP mice (Figure 6B-ii).

**Figure 5.**
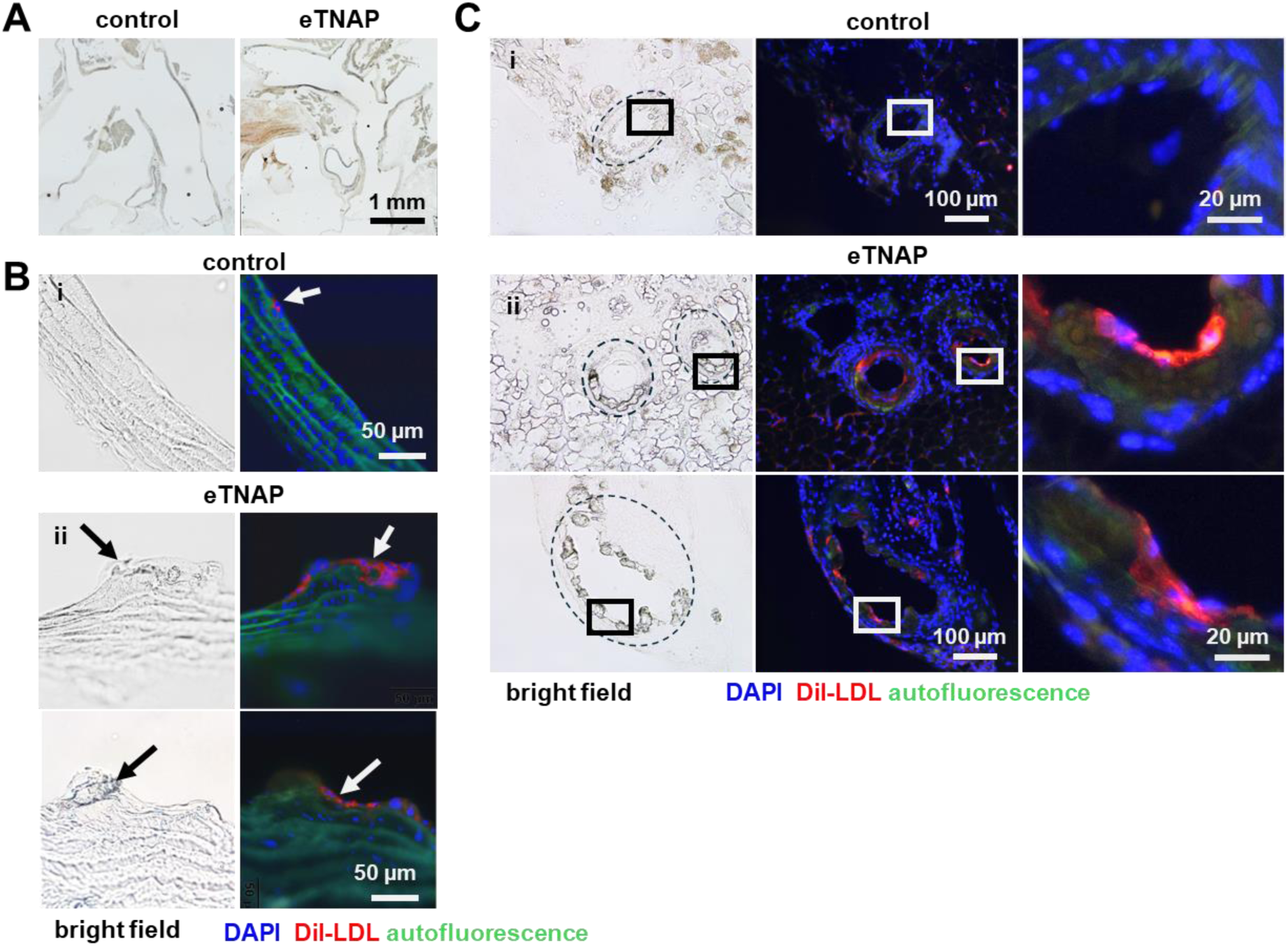
Effect of subendothelial calcification on native LDL uptake *in vivo*. A. Longitudinal sections of the aortic arch from control and eTNAP mice; B. (i) representative example of native LDL uptake in control mouse aortic arch (white arrow), (ii) representative examples of LDL uptake in the aortic arch of eTNAP mice; black arrows point to the irregularities of the aortic surface, white arrow point to the LDL in the same lesion; C. LDL uptake in mesenteric arteries of (i) a control mouse and (ii) eTNAP mice; boxed areas are enlarged (right panels); blue, nuclear stain (DAPI); red, fluorescent LDL (DiI-LDL); green, autofluorescence.

**Figure 6.**
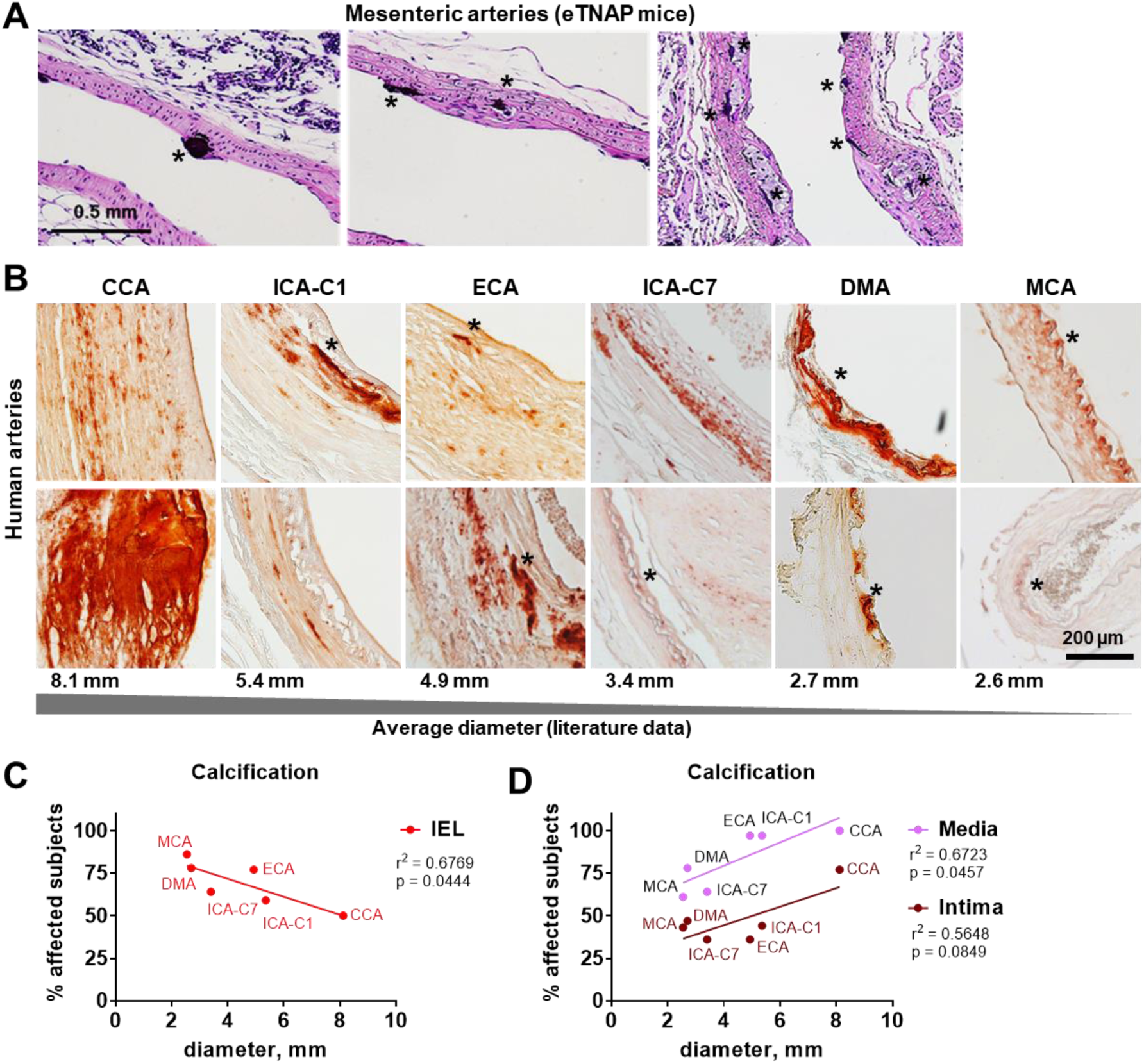
Internal elastic lamina (IEL) calcification in eTNAP mice and human arteries. A. Subendothelial calcification in the mesenteric arteries of eTNAP mice; *, IEL calcifications detected by hematoxylin (deep blue); B. Detection of calcification in human medium-sized arteries by alizarin red staining; CCA, common carotid artery; ICA-C1, the internal carotid artery extracranial C1 segment; ECA, external carotid artery; ICA-C7, distal C7 segment of the ICA; DMA, Drummond marginal artery of the colon; MCA, middle cerebral artery; *, IEL calcifications; average diameters of the arteries were cited from the literature;^32–34^ C. Negative correlation between arterial diameter and the prevalence of IEL calcification (% affected) in human subjects; D. Positive correlations between arterial diameter and medial/intimal calcification in human subjects.

### IEL calcification in human medium-sized arteries

To address the potential relevance of our observations to human vascular disease, we investigated the frequency of IEL (internal elastic lamina) calcification in vascular samples obtained from cadavers. We focused on IEL calcification because of its similarity with subendothelial calcification in our eTNAP model, in which the IEL is prominently involved (Figure 6A). Non-decalcified samples were cryosectioned and stained with alizarin red to detect calcification in the IEL, intima, and media (Figure 6B). We found the highest prevalence of IEL calcification in the middle cerebral arteries (MCA, 86% affected subjects), followed by the Drummond marginal artery of the colon (DMA, 78%) and the external carotid artery (ECA, 77%, Table 2). Interestingly, we found a negative correlation between the prevalence of IEL calcification and the average diameter of the arteries (obtained from the literature,^32–34^ p = 0.0444, Figure 6C). In contrast, the prevalence of intimal and medial calcification showed a positive trend with arterial diameter, with the highest degree of media and intima calcification found in the common carotid arteries (Figure 6D, Table 2). The distinct relationship between IEL calcification and arterial diameter may suggest that the pathophysiology of IEL calcification differs from that of more recognized medial and intimal calcification.

**Table 2.**
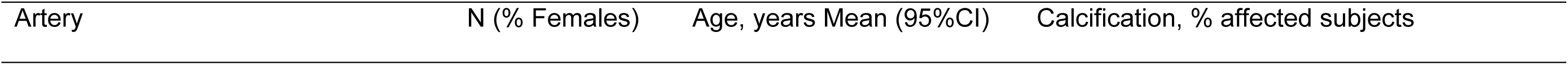
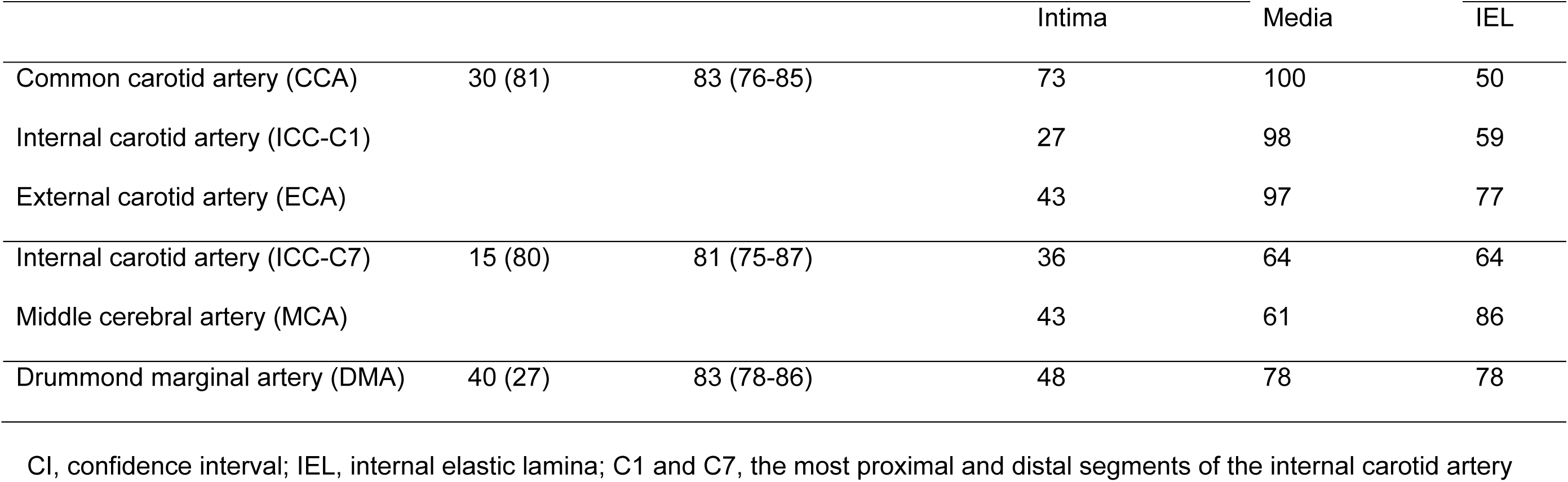
Calcification in the intima, media, and internal elastic lamina detected in human arteries by alizarin red staining Artery N (% Females) Age, years Mean (95%CI) Calcification, % affected subjects.

|  |  |  | Intima | Media | IEL |
| --- | --- | --- | --- | --- | --- |
| Common carotid artery (CCA) | 30 (81) | 83 (76-85) | 73 | 100 | 50 |
| Internal carotid artery (ICC-C1) |  |  | 27 | 98 | 59 |
| External carotid artery (ECA) |  |  | 43 | 97 | 77 |
| Internal carotid artery (ICC-C7) | 15 (80) | 81 (75-87) | 36 | 64 | 64 |
| Middle cerebral artery (MCA) |  |  | 43 | 61 | 86 |
| Drummond marginal artery (DMA) | 40 (27) | 83 (78-86) | 48 | 78 | 78 |

CI, confidence interval; IEL, internal elastic lamina; C1 and C7, the most proximal and distal segments of the internal carotid artery

## Discussion

Vascular calcification and conditions predisposing to vascular calcification, including diabetes and chronic kidney disease, confer a higher risk for ASCVD.^4–8^ Historically, calcification has been interpreted as a consequence of atherosclerosis or arteriosclerosis, depending on which arterial layer (intimal or medial) was affected.^35, 36^ Our studies demonstrated that calcification, induced by endothelial TNAP expression, increases atherosclerosis, while pharmacologic inhibition of TNAP activity reduces the natural progression of atherosclerosis in mice.^25, 37^

In this study, we tested the hypothesis that subendothelial calcification, such as that affecting the IEL, can initiate atherosclerosis by increasing endothelial roughness and disturbing WSS, thus increasing LDL uptake and plaque formation. Our mouse model confirmed that endothelial overexpression of TNAP, an enzyme that degrades the calcification inhibitor PPi (pyrophosphate), leads to subendothelial calcifications and accelerates atherosclerosis independent of the levels of plasma lipids (Figure 1). Subendothelial calcification increases surface roughness (Figure 2, Table 1) and leads to uneven redistributing WSS on the endothelium (Figure 3). Low-shear stress pockets associated with subendothelial microcalcifications are likely responsible for an increase in LDL uptake by endothelial cells, as shown by our *in vitro* and *in vivo* studies (Figures 4 and 5). Although an intriguing discovery, the exact phenotypic changes in endothelial cells in response to subendothelial calcification were outside the scope of our current study, and their molecular mechanisms^38^ remain a subject of further investigation.

We also found that IEL calcification was prevalent in an elderly human donor population. IEL calcification was more prevalent in smaller-sized arteries, whereas intimal and medial calcification correlated positively with larger vessel diameters, thus suggesting a distinct mechanism of IEL calcification (Figure 6). Whether IEL calcification is an early pathologic change preceding human atherosclerosis cannot be discerned from our study.

Vascular calcification, a strong predictor of ASCVD,^4–8^ currently has no therapeutic strategies except for the invasive intravascular ablation of severely calcified atherosclerotic lesions in symptomatic patients.^39, 40^ New therapeutic targets, including TNAP inhibition, are being developed.^41^ TNAP activity in the blood (measured b a plasma alkaline phosphatase [ALP] test) is positively associated with CVD and all-cause mortality.^42^ Our study offers a potential mechanism linking ALP activity and ASCVD, supporting the validity of TNAP as a therapeutic target for vascular calcification.

## Conclusions

Subendothelial calcification increases arterial lumen roughness, perturbs near-wall hemodynamics and promotes lipid deposition. W suggest that IEL calcification may initiate atherosclerosis. Further studies should examine the validity of this causative relationship in human atherosclerosis.

## Limitations

The major limitation of this study is the lack of epidemiologic evidence supporting the relationship between subendothelial calcification and the onset of human atherosclerosis. Therefore, our theory that certain forms of calcifications, such as IEL calcification, can initiate atherosclerosis should promote future research.

## Acknowledgments

This work was supported by the National Institutes of Health grant R01HL149864. The NYITCOM DO/PhD program supported KAB; the NYITCOM Academic Medicine Scholar Program supported NDK and AM; DP, NV, and JD received funds from the NYITCOM Summer Research program. Micro-CT studies were conducted at the NYIT Visualization Center (NSF grant 1828305).

